# Unlocking precision: How corneal cell area analysis revolutionizes post-transplant stem cell monitoring

**DOI:** 10.1101/2024.09.17.612429

**Authors:** Patrick Parkinson, Irina Makarenko, Oliver J Baylis, Gustavo S Figueiredo, Majlinda Lako, Anvar Shukurov, Francisco C Figueiredo, Laura E Wadkin

## Abstract

The corneal epithelium is maintained by limbal stem cells (LSCs). Dysfunction of the LSCs, resulting from chemical and thermal burns, contact lens-related disease, congenial disorders, among other conditions, leads to limbal stem cell deficiency (LSCD), a sight-threatening condition. An effective treatment of LSCD, with 76% of patients reporting regained sight up to 24 months after the operation, consists of transplanting *ex-vivo* cultured LSCs from the patient’s other healthy eye (i.e. autologous) or donor (i.e. allogeneic) to the affected eye. The post-operative assessment of corneal recovery is crucial but relies on ponderous and generally subjective visual inspection of a large number of microscopic images of the corneal epithelial cells, relying on the personal experience of the practitioner to interpret imprecise, qualitative diagnostic criteria. From a unique library of 100,000 cornea cell images from 34 patients, we have randomly selected 10 individuals (3,668 images) to demonstrate that the frequency distribution of the epithelial cell areas is a sensitive diagnostic tool of the corneal epithelium status. After a successful operation the distribution of cell areas is rather flat, reflecting an anomalously wide range of cell areas. As the cornea recovers, the frequency distribution becomes narrower with high statistical confidence and eventually approaches that of the healthy cornea. The corneal epithelial cell shape is independent of the cornea status despite a widespread expectation that healthy cells have a hexagonal shape. We also show that the corneal epithelial cell area distribution and its variation with the depth within the cornea are specific to each patient.

**Significance Statement:** Chemical and thermal cornea burns, contact-lens damage and hereditary factors, among other conditions, cause limbal stem cell deficiency (LSCD), a widespread sight-threatening condition. An efficient LSCD treatment involves a stem cell transplant from the patient’s other healthy eye, in unilateral cases, or a donor, in bilateral cases. Traditional post-operative cornea monitoring is laborious and often subjective as it relies on visual inspection of microscopy corneal epithelial images. We show that the distribution of epithelial cell areas is a sensitive LSCD diagnostic, evolving systematically to a healthy form after a successful treatment. We have developed computer algorithms to implement this quantitative, sensitive and precise approach which can radically improve the quality of both cornea monitoring in disease and response to treatment.

## Introduction

The cornea is a transparent structure forming the front layer of the eye with the corneal epithelium comprising the outer 50—60μm (1—3). The corneal epithelium is stratified into five to seven cellular layers consisting of three distinct cell types (2—4). Comprising the outermost two to three layers of the epithelium are the flat, squamous (poorly defined) superficial cells with a large surface area and an approximate thickness of 2—6μm. These are followed by two to three layers of the more structurally defined wing cells, each with an approximate thickness of 5--10μm. At the base of the epithelium is a monolayer of columnar basal cells. These cells are approximately 20μm tall with a considerably smaller surface area than the other cell types. They are the only cells in the corneal epithelium capable of mitosis and are responsible for the generation of the other cell types as transient amplifying cells (TAC).

The corneal epithelium is maintained by the limbal stem cells (LSCs) (5). Damage, loss or dysfunction of the LSCs results in a disease known as limbal stem cell deficiency (LSCD) (5). LSCD is a sight-threatening condition resulting from damage to the limbus, the region between the cornea, conjunctiva and the sclera where LSCs reside, that is responsible for the repair and renewal of the corneal epithelium (6,7). Damage to the limbus has a number of possible causes including chemical and thermal burns, contact lens-related disease, hereditary disorders (such as aniridia and ectodermal dysplasia) and inflammatory disease (such as Stevens-Johnson syndrome and ocular cicatricial pemphigoid) (5).

With the onset of LSCD, the normal process of homeostasis cannot occur and the barrier action of the limbus between the cornea, conjunctiva and the sclera is compromised. A process of cornea neovascularisation and conjunctivalisation occurs, whereby the opaque conjunctival epithelial cells migrate into the cornea replacing the transparent corneal epithelial cells. This may result in recurrent epithelial erosions, persistent epithelial defects, and neovascularization of the superficial cornea. Common presentations are chronic inflammation, with red eyes and scarring as well as loss of vision (8), with severe long-term consequences for the patients’ quality of life, their families and society (9). The reported incidence of ocular chemical injuries in the UK ranges from 7—10% among all ocular (10). Burns occur most frequently in and around industrial areas, at patient’s homes, or as a result of criminal assault (acid attacks). The number of attacks in the UK has increased significantly in previous years, with assaults more than doubling in the period 2012 to 2016 (10).

Treatment for patients with total LSCD comes in the form of a stem cell transplant (8, 11, 12). This begins with the *ex-vivo* culturing of limbal epithelial grafts from small biopsies taken from a donor eye (8,13). This can be the patient’s other healthy eye (in the case of unilateral LSCD), or a cadaver donor (in the case of bilateral LSCD). After undergoing *ex-vivo* expansion on a human amniotic membrane (HAM) matrix, the autologous limbal epithelium is then transplanted onto the affected cornea, after a superficial keratectomy to remove the conjunctivalised scarred tissue (8). The subsequent proliferation and differentiation of LSCs across the cornea reverses the conjunctivalisation process. The procedure has been shown to accelerate the recovery of the corneal surface, encourage the closure of epithelial defects, eliminate or decrease corneal vascularity and consequently improve the visual acuity (8). The treatment has proven to be very successful, with ∼76% visually impaired patients regaining sight on a timescale of 24 months (14).

The diagnosis and post-operative monitoring of the LSCD corneas relies on the visual inspection of cellular images, usually obtained using *in-vivo* confocal microscopy (IVCM), a laborious and often subjective means of investigation (5). Furthermore, the differences between the healthy, damaged, pre- and post-operative corneal epithelial structures are subtle and not easily discernible (15). Computer-aided, quantitative analysis of such images is therefore needed for making objective unbiased decisions on the success of LSC transplantation (16).

In this study, we quantitatively analyse IVCM corneal epithelium images of patients with total LSCD (8). The data consist of over 100,000 images from 34 LSCD patients taken both before and (up to two years) after LSC transplant. Images are taken across the entire thickness of the corneal epithelium (0--60μm) in each of the central and four peripheral corneal regions (Fig. 1A). All patients had total, unilateral LSCD resulting from a chemical burn and received a unilateral, autologous LSC transplant. For each patient, we also have similarly obtained images of the healthy corneal epithelium from the contralateral, unaffected eye. These images were collected, over the last decade, by a small group of clinicians and are therefore part of a uniquely complete, homogeneous, unbiased and representative database with respect to patient diagnosis, treatment and the systematic and stable data collection protocols. The analysis carried out in this study focuses on a sample from ten patients, consisting of 3,688 of these images to first develop and refine the imaging strategy and image analysis tools before increasing the size of the data set (17).

**Figure 1.**
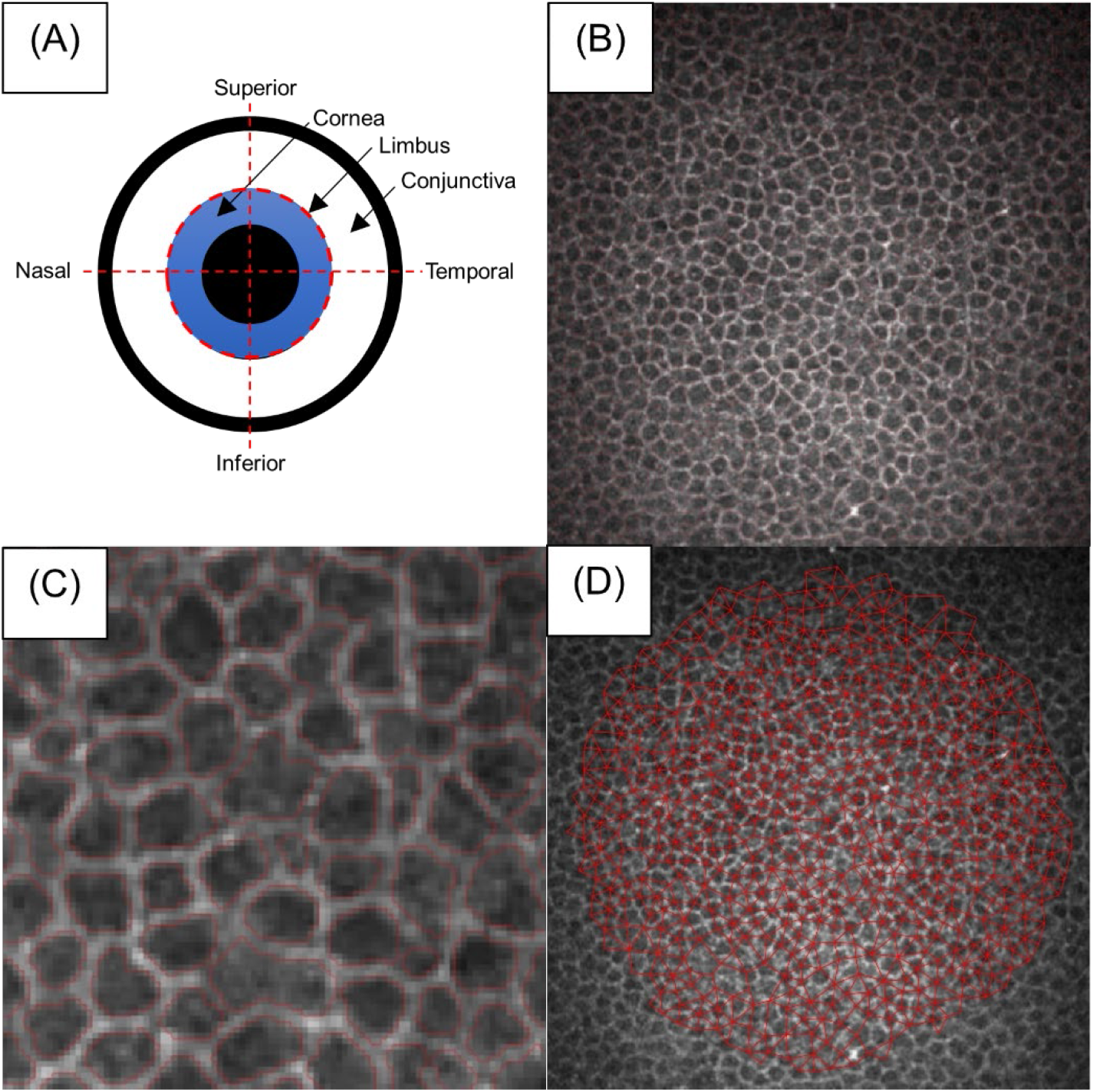
An example of the identification of cell boundaries using the segmentation algorithm detailed in **Materials and Methods**. (A) Location of the limbus and the inferior, nasal, superior and temporal regions of the corneal epithelium (20). (B) Original IVCM image (4.3 × 10^−11^ *μm*^2^) of the central corneal epithelium at a nominal depth of 23μm in a healthy eye. (C) The cell boundaries identified in the central part of the image at 1/16 scale. (D) Voronoi tessellation, with Gabriel graph overlaid, for the same area as in Panel (B). An average of 650 corneal epithelial cells were detected and segmented in each IVCM image.

We have developed an algorithm for the identification of cell boundaries in the IVCM images (Fig. 1) which performs better than existing algorithms, such as that in the widely used *ImageJ* software package (18). Extracting morphological data regarding individual cells, we identify the probability distribution function (PDF) of the epithelial cell areas as a sensitive diagnostic of the state of the cornea. The diagnostic relevance of the average epithelial cell area has been suggested earlier (19) but the PDF, representing the fraction of the number of cells with a given area out of the whole range of cell sizes, offers a by far more detailed, sensitive and reliable measure of the state of the epithelium, from which the mean cell area can easily be derived. On the other hand, the shape of the corneal epithelial cells, a diagnostic criterion often used in practice, does not appear to vary with corneal health, suggesting it is of no diagnostic value. We also analyse images of the healthy contralateral cornea to characterise its structure and assess variation between individual patients, investigating the existence of the notion of a well-defined, universal cell structure in the epithelium of a healthy cornea, which could be used as a reference for the disease affected.

## Results

### Cell area distribution

The analysis of the IVCM images starts with the identification of the cell boundaries in each image, discussed above and in **Materials and Methods**, as illustrated in Fig.1. Utilising the output, we derive the PDF of the cell areas, in practice, a histogram of the cell areas normalised to the unit area under the histogram (so that images with different total cell numbers are directly comparable). The value of the PDF represents the fraction of cells of a given area.

PDF analyses were performed for the LSCD affected corneal epithelium as well as the contralateral healthy corneal epithelium, in the central and four peripheral corneal regions (Fig. 2). In all regions, including for other patients (see Figs. **S1--S3** in **SI Appendix**), we found that the distribution of cell areas is not only responsive to LSCD, but recovers a healthy distribution over a 24-month period post-transplant. The PDFs for the normal eye have a pronounced maximum and a weak tail at large areas: the variation in cell size is relatively small and they have a well-defined typical size in all cases. At six months after the LSC transplantation, there is a large number of cells with anomalously large and small areas, the PDFs do not have a sharp single maximum, and in some cases (Fig. 2A) even have two local maxima. Remarkably, the fractions of both the smaller and, especially, larger cells gradually decrease as the PDFs approach those of the healthy eye 24 months after the transplantation. These data support the notion that the corneal epithelial cell area distribution is sensitive to damage similarly in the centre and all four peripheral regions of the cornea. It suggests probability density of cell areas is useful as a tool for monitoring the recovery of the entire LSCD corneal epithelium layer post-treatment.

**Figure 2.**
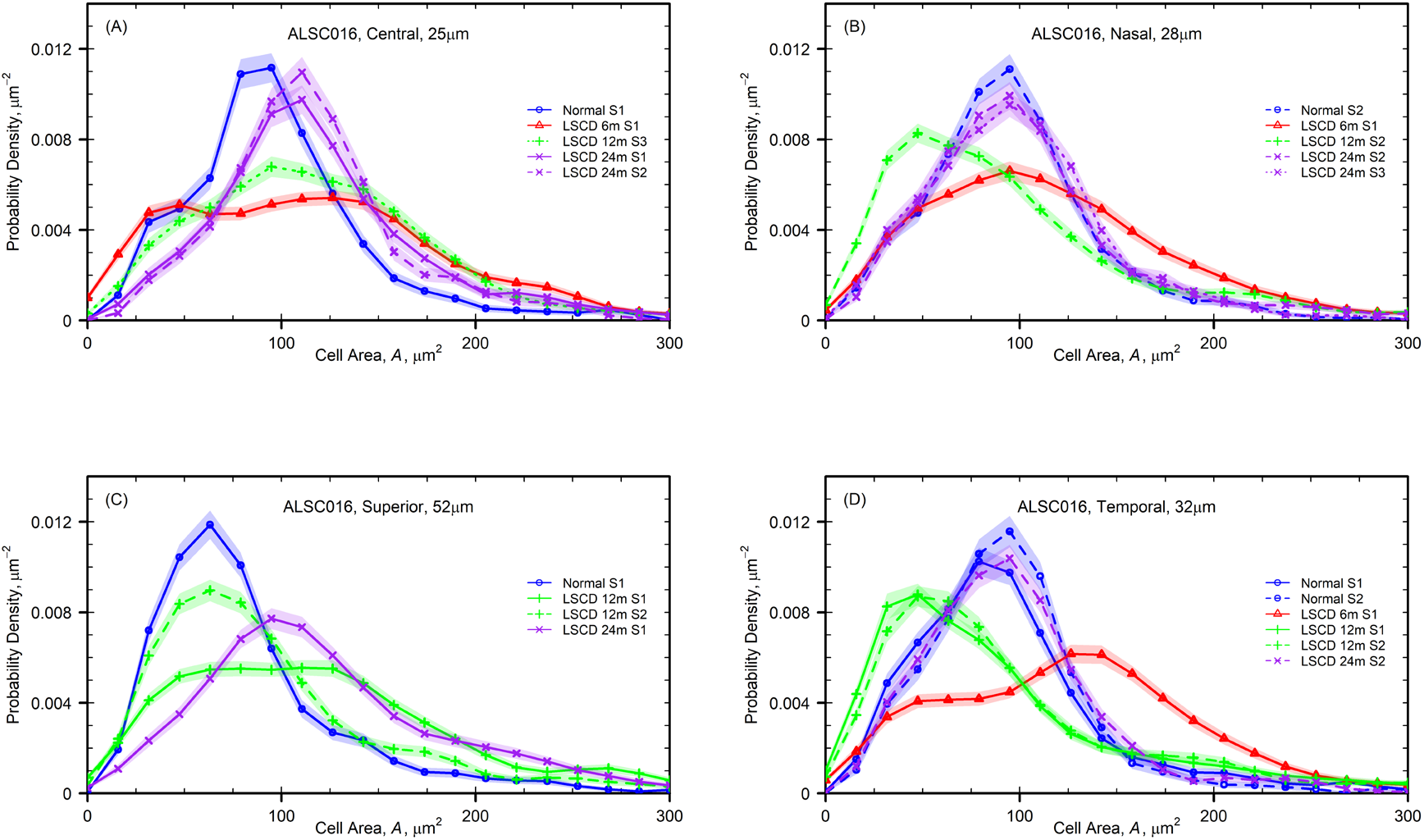
The PDF of cell areas of patient ALSC016 in (A) the central corneal region at a depth of 25μm, (B) the nasal corneal region at a depth of 28μm, (C) the superior corneal region at a depth of 52μm (D) the temporal corneal region at a depth of 32μm, for the healthy (blue lines/circles) and post-operative central cornea at six months (red lines/triangles), 12 months (green lines/plus signs), and 24 months (purple lines/crosses) after surgery. The PDF for the normal eye has a pronounced maximum in all cases (at 87μm^2^, 91μm^2^, 65μm^2^ and 96μm^2^ (S2), in Panels(A)--(D), respectively) and a weak tail at large areas. The statistical difference of the PDFs at six, 12 and 24 months from the healthy distribution are assessed by the Kolmogorov-Smirnov test at the 1% significance level (see Fig. **S5** in **SI Appendix**). The shaded regions show the bootstrap 2σ errors (see **Materials and Methods** for details). Labels S1--S3 indicate from what sequence of images these were taken, with PDFs differentiated by line type.

### Variation of cell area with depth

The stratification of the epithelium by different cell types (as discussed in the **Introduction**) should result in a systematic trend of a decreasing cell area with depth (17, 21). Here we explore the relationship between cell size and depth in order to, firstly, assess its variation between individuals and, secondly, its potential as a diagnostic feature of the cornea health.

The variation of the cells area with depth within a healthy cornea is discussed by Sterenczak et al. (17) who consider the average cell area at a given depth for 25 individuals, performing a single fit to all. Our data demonstrate that the rate of change of the cell area with depth differs between individuals so strongly (Fig. 3) that drawing conclusions from a fit to all cases may be misleading: in other words, it is not possible to introduce the notion of a `standard’ healthy cornea which might be used as a reference when assessing its health. Importantly, we note that the PDFs of the cell area at a fixed depth differ significantly between individuals (see Figs. **S1—S3** in **SI Appendix**), further evidence to this point.

**Figure 3.**
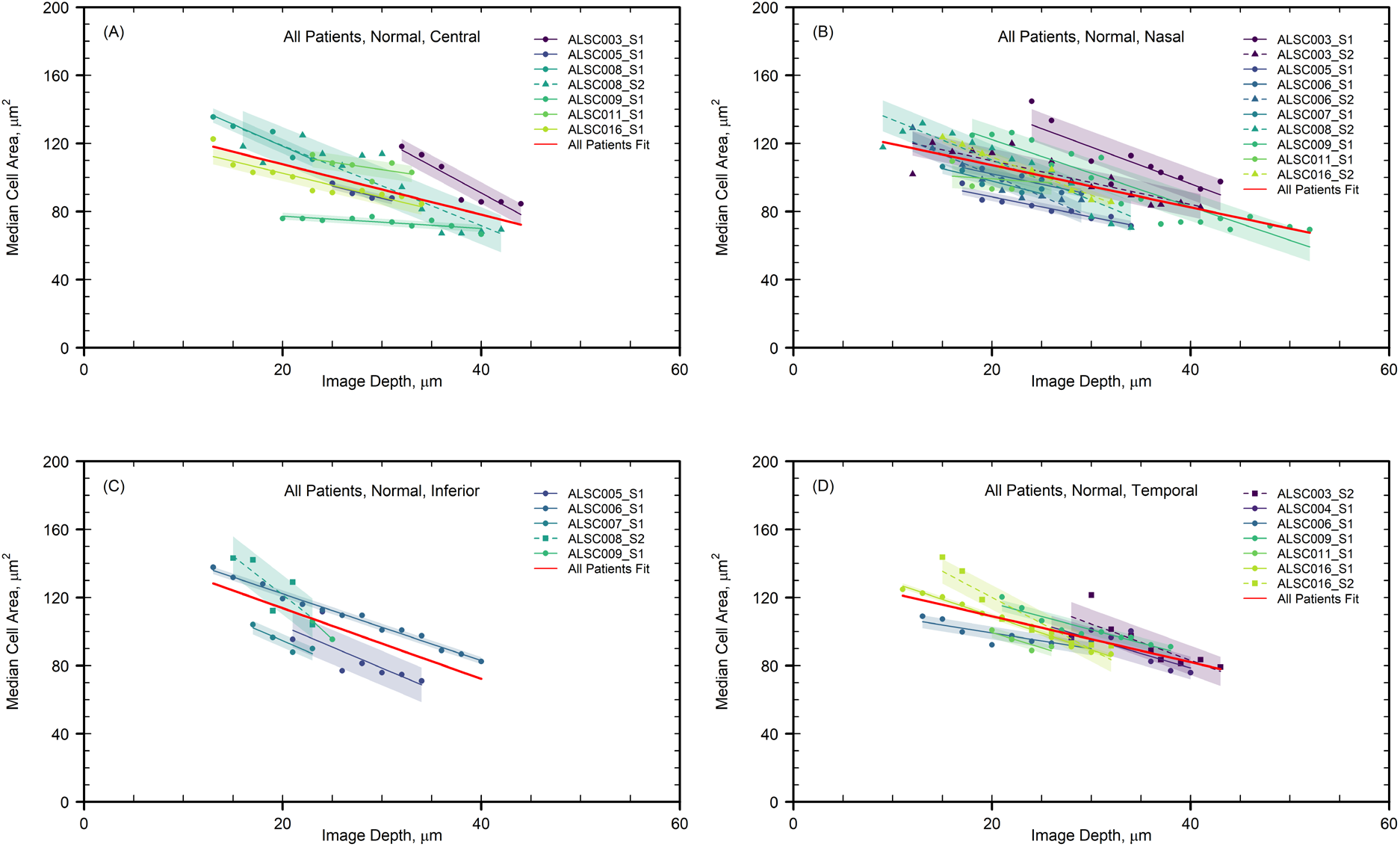
Distribution of median cell areas varying with depth for all patients in the healthy corneal epithelium, in the (A) central, (B) nasal, (C) inferior and (D) temporal regions. The coloured points are the individual medians at a given depth, for a given patient, with different patients distinguished by colour and different sequences distinguished by point shape. Red lines depict the linear relationship between depth and cell area, given by equation (1). This relationship was derived by performing a linear regression fit to all data points plotted. All other lines depict similar fits for each patient separately. The corresponding shaded regions show the 2σ uncertainty of the coefficient of depth for each fit. It is clear that all coefficients are negative. Depth is significant in the modelling of cell area in all fits at the 1% level (see Table **S1** in **SI Appendix**).

Plotting median cell areas as a function of depth for all sequences of healthy corneal epithelium images in each of the corneal regions (Figs. 3 and **S6**) we fit to these data points, with high statistical significance, a linear relation between the depth *D* and cell area *A* of the form

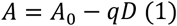

where the coefficients *A*_*0*_ and *q* are obtained using the least-squares fit to all the patient data for each region of the cornea (see Table **S1** in **SI Appendix**). It is clear that all coefficients are negative. Depth is significant in the modelling of cell area in all fits at the 1% level. Cell areas systematically decrease with the depth but different individuals have different fit parameters *A*_*0*_ and *q*, and this difference is statistically significant. We present in Table **S2** in **SI Appendix** tables of fit parameters for each individual patient in the central cornea to confirm this conclusion.

In order to find out if the cell area variation with the depth within the cornea can be used as a diagnostic indicator of corneal health, we perform the same analysis for LSCD affected corneal epithelium images (see Fig. **S7** in **SI Appendix**), with fits both to all patients and individuals, at various times after the transplant, in the central and inferior corneal epithelium (see Table **S3** in **SI Appendix**). It appears that the values of both *A*_*0*_ and *q* obtained from the all the data available (without distinguishing individual patients) are similar between the healthy cornea (the first and fifth rows) and 24 months after the operation (the fourth and eight rows) but not in the early post-operative period (the second, third, sixth and seventh rows). This may suggest that the variation of the median cell area with the depth can be an appropriate additional diagnostic criterion, independent of the cell area PDFs discussed above. This assertion needs to be carefully tested using the data of individual patients and a larger data set.

### Cell shape

A roughly hexagonal shape has been suggested as a diagnostic feature of healthy wing cells in the corneal epithelium (22), but the validity of this diagnostic has been questioned in some studies (17). We hypothesised that cell shapes could be another informative, and crucial, parameter of post-operative cornea monitoring.

In order to quantify a cells shape, we identified the number of nearest neighbours of the cells (for example, hexagonal cells would have six neighbours) in segmented IVCM images (see **Materials and Methods**). Performing PDF analyses once again, we find that the most probable number of neighbours is five, so the cells have a roughly pentagonal shape (Fig. 4). We found this to be consistent across all four regions in the peripheral cornea and across all depths in every case, for every patient. Moreover, the cell shape is independent of the state of the cornea: the distributions for the post-operative eye at six, 12 and 24 months are similar to that of the normal cornea. Corneal epithelial cells have a roughly pentagonal shape in both the healthy and LSCD affected epithelium.

**Figure 4.**
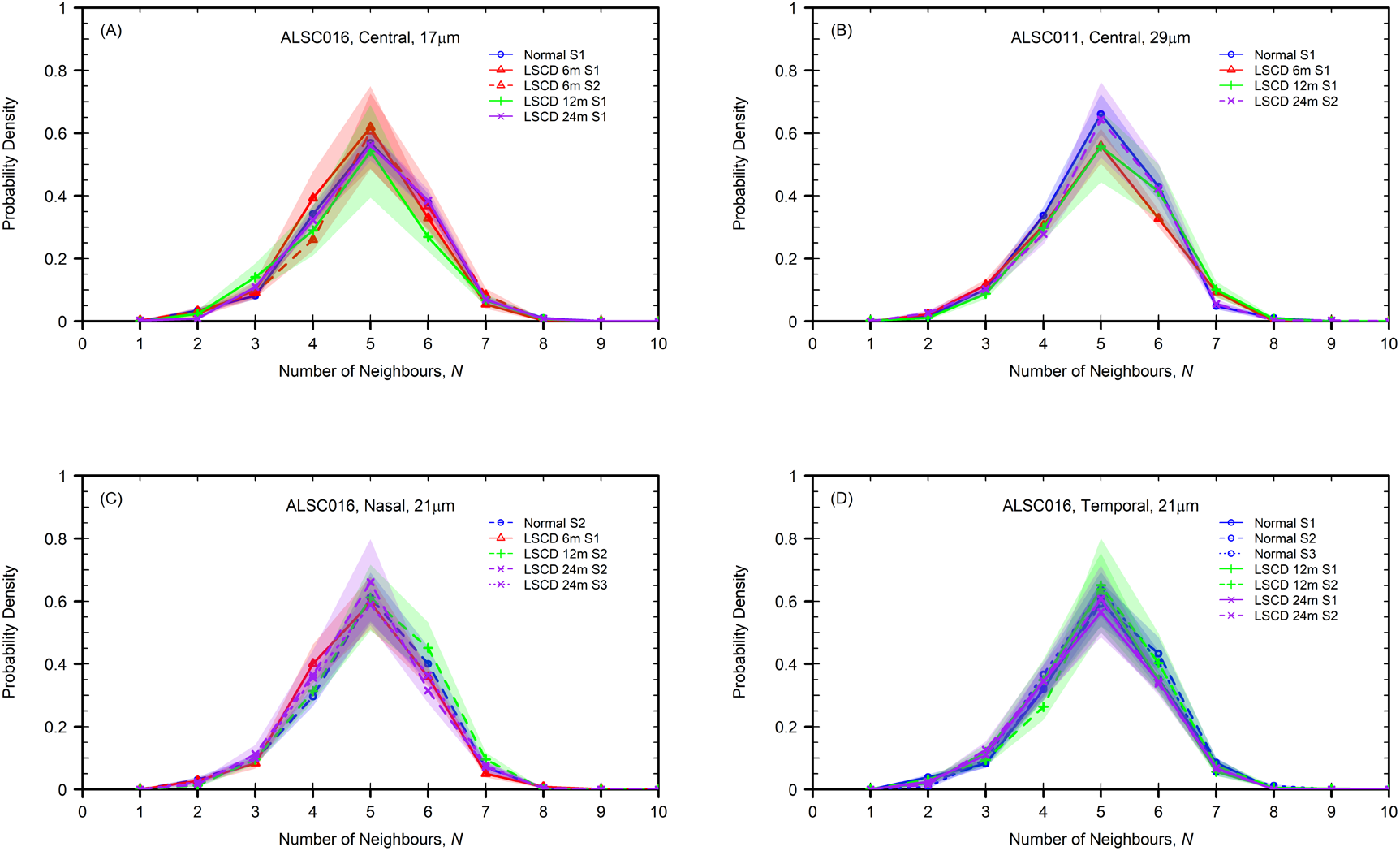
Distributions of the number of neighbours of corneal epithelial cells in (A) the central cornea region of patient ALSC016 at a depth of 17μm (B) the central cornea region for patient ALSC011 at a depth of 29μm, (C) the nasal cornea region for patient ALSC016 at a depth of 21μm and (D) the temporal cornea region for patient ALSC016 at a depth of 21μm, for the healthy (blue lines/circles) and post-operative central cornea at six months (red lines/triangles), 12 months (green lines/plus signs), and 24 months (purple lines/crosses) after surgery. The statistical similarity of the PDFs are confirmed using the two-tailed Kolmogorov--Smirnov test with a significance level of 1% (see Figs. **S8** and **S9** in **SI Appendix**). The shaded regions show the bootstrap 2σ errors (see **Materials and Methods** for details). Labels S1–-S3 indicate from what sequence of images these were taken, with PDFs differentiated by line type. It is clear in all cases that most cells have five neighbours.

## Discussion

In this study we analysed several morphological features of corneal epithelium cells (mainly, the wing cells) which can serve as indicators of corneal epithelial health. We show that cell area not only varies between the healthy and LSCD-affected eye, but that it is responsive to post-treatment recovery, representing a sensitive diagnostic of the cornea damage or health. Meanwhile, the cells shape remains consistent throughout the healthy, diseased and recovering cornea and so is unlikely to be useful for diagnostic purposes. We suggest the probability distribution function (PDF), the fraction of cells of a given area evaluated for the whole range of areas, as an appropriate characteristic of the cell area. It has been suggested prior to this analysis that mean basal cell size could provide a useful diagnostic of the state of the cornea (19). The mean area is just one of the parameters of the PDF, so the PDF as a whole represents a significantly more sensitive and informative diagnostic.

It is the uniqueness of our data set that motivated our initial investigations. Much of the existing literature regarding the corneal epithelium concerns image analysis of IVCM images of the central region (17, 21). Our image library covers the entire corneal epithelium, including the peripheral regions. With documented variation in epithelial structure across these regions, both in layered structure with depth (4) and the presence of other anatomical features, for example the Palisades of Vogt (1), it was clear to us that conclusions made about the central corneal epithelium may not necessarily be generalisable to the entire epithelial layer.

Our approach relies on the comparison of the cell morphology in the healthy contralateral eye of a patient and in the recovering, post-operative cornea and thus only applies in the case of the unilateral LSCD. Therefore, we explored the possibility of a universal, generally applicable description of the epithelial cell morphology in a healthy cornea. Our results indicate that different individuals have statistically different morphological cell parameters, so the notion of a `standard healthy cornea’ may not be pertinent. On the other hand, the fact that the morphology of the corneal epithelial cells 24 months after a successful operation becomes very similar (if not identical) to that in the healthy contralateral eye of the *same patient*, suggests strongly that the healthy eye of the same person can be used as a reference when assessing the state of the other damaged eye (see also (23)).

We have found some evidence that the variation of the median cell area with the depth within the epithelium may be another useful diagnostic tool of the cornea damage. However, we show that the cell sizes and the rate of their change with the depth vary rather widely from one individual to another. Therefore, the cell area variation assessed across a number of individuals (17, 21) does not appear to be useful for the diagnostic purposes: each individual has to be considered separately. The choice to use the median cell area in our study as opposed to the mean value (as in (17)) was motivated by significant variation observed in cell areas extracted from the IVCM images. In some cases, there is an abundance of unusually large cells (evident in the cell area PDFs) which have a disproportionate effect on the mean cell area but not so much on the median. Such large cells appear widespread in the damaged cornea but could also be an artefact of imperfect segmentation when an image is not of sufficient quality to resolve cell boundaries at all positions in the image. As a result, the uncertainty of the mean cell area is 2--3 times larger than that of the median. It has been observed, in some cases, that when the segmentation algorithm is unable to identify cell centres multiple corneal cells can be segmented as one, effectively producing a cell with a much greater area. Such a value distorts the mean cell area of all the cells in an IVCM image, but not the median. Even if this is a nuance of our segmentation algorithm alone, the median is known to be a more stable diagnostic than the mean in many similar contexts. Our most reliable results are obtained for the central region of the corneal image, where increased brightness and image sharpness make for reliable segmentation. We find that the segmentation is less reliable close to the boundary of the central region and in the peripheral cornea parts.

Unlike previous studies, we have images of both the healthy cornea and the contralateral LSCD-affected eye of the same individual, a unique feature of our IVCM data set. By performing the same analysis on our LSCD-affected images, we were able to show that the above conclusions hold for damaged eyes also, at all stages of recovery. Though cell size is affected by LSCD, the individual cell types remain stratified, and thus the trend of decreasing cell area with depth remains.

Each IVCM image has the depth label measured against the reference value, hand-entered by the operator near the cornea surface at the start of each image sequence. Because of the unavoidably subjective and rather unsystematic nature of the reference point, the recorded depth is often not accurate enough to identify images taken at the same true depth in different sequences. Since the cell parameters vary with the depth, this makes image comparison more difficult. We have applied a significant effort to verify and correct the recorded depths as to ensure reliable comparisons of different images.

Similarly to other existing algorithms (17), our image segmentation algorithm, in its present form, most effectively segments wing cells. Due to the phenotypic differences between cell types, any adjustment of segmentation parameters to improve segmentation of one cell type would affect the segmentation efficiency for another. This is why most of the images suitable for good segmentation are within the depth range 15-30μm, where the wing cells are located. However, the basal cells are of particular interest to study further, being the only cell type capable of mitosis (24).

Regarding the cell shape, our analysis shows that epithelial cells typically have five neighbours in the healthy cornea. This is consistent with the findings of Sterenczak et al. (17), however we found no depth dependence as they report. Our findings are consistent in all regions of the cornea, at all depths. Further to this we find that the cells shape (characterised by the number of neighbours) is independent of the health of the cornea. Sterenczak et al. (17) find that the number of cell neighbours decreases from the anterior to the posterior epithelium. This may be explained by the change in cell structure as we move into the basal cell layer.

Our results rely on the parameters of the wing cells but other cornea cell types might also provide useful diagnostics of LSCD and other conditions. Furthermore, there are contradictory results regarding the dependence of various cornea cell parameters on the age, gender and demographic factors (17, 22, 23, 25, 26). These directions deserve further analysis, and the extensive library of IVCM images available to us offers a unique opportunity for that.

## Materials and Methods

### Study design

A 3-year, phase II, prospective, single centre, interventional study was conducted in accordance with the principles of the Declaration of Helsinki and approval was obtained from the local Research Ethics Committee (11/NE/0236), MHRA Clinical Trial Authorization (17136/0254/001-0001) and under an HTA licence (11122). The clinical trial was also registered with the European Union Drug Regulating Authorities Clinical Trials Database (EuDRACT 2011— 000608-16) and the ISRCTN (International Standard Randomised Controlled Trial Number (ISRCTN51772481). All patients provided written informed consent.

Consecutive patients presenting with acquired and unilateral LSCD due, in the great majority, to ocular surface burns (i.e., chemical and thermal) at the Department of Ophthalmology at the Royal Victoria Infirmary, Newcastle upon Tyne Hospitals NHS Foundation Trust, Newcastle upon Tyne, UK were invited to take part in the clinical trial and subsequently enrolled in the study. Patients were recruited between June 2012 and January 2015 and followed-up over a 36-month period following autologous corneal limbal epithelial transplantation (CLET).

### IVCM images

The cornea images were collected using *in-vivo* confocal microscopy (IVCM - HRT3, Heidelberg, Germany). The IVCM technique is well described in previous papers (27). The image collection process was similar in both LSCD-affected and healthy eyes. Each raw image is accompanied by an image frame containing relevant meta-data, including a depth label, OS/OD (left/right) identification and the date the image was taken.

For each patient, we have some or all of the following: images of the centre and four peripheral regions of the normal (healthy) cornea, images of the contralateral LSCD-affected eye pre-transplant (baseline images) and images of the cornea at six, 12, 18, 24 and 36 months post-LSC transplant. Not every patient has images covering all these post-operative times and not all images are suitable for quantitative analysis. For example, the baseline images exhibit significant damage and therefore very little or no identifiable cell structure (see Fig. **S10A** in **SI Appendix**). Such images are not used in our analysis. There are also images taken outside the depth range of the corneal epithelium (approx. 0--60μm, see Fig. **S10B** in **SI Appendix**). These images are discarded too. There are also poor-quality images of the corneal epithelium which are not suitable for the analysis as they are affected by the motion of the patient’s eye. Images of the four peripheral cornea regions suffer from this more often than the central region.

Following removal of the unsuitable images, we are left with an analytic database of 1,483 images, covering a 0--60μm depth range within the anterior cornea, taken at all five locations within healthy, baseline (damaged) and post-operative (recovering) cornea. Image availability varies from patient to patient.

### Morphological Segmentation

- In order to quantify characteristics of IVCM images at a single-cell level, we initially used the morphological segmentation feature in the open-source image processing package Fiji, employing an ImageJ segmentation algorithm (see Fig. **S11** in **SI Appendix**). However, it soon became clear that subtle differences in the cell morphology between the central and peripheral cornea regions (4) could not be captured with ImageJ. Therefore, we developed an alternative segmentation algorithm in MATLAB described below (Fig. 1B--C). With a significant improvement in resolution, this algorithm is capable of detecting the slight morphological differences between the central and peripheral cornea. Following the segmentation, the following morphometric cell features are calculated: the cell area, aspect ratio (AR), circularity, hexagonality and number of neighbours. These metrics are output in a file with a separate entry for each segmented cell in the image. The algorithm also employs Voronoi tessellation to identify cell centres which can be used to extract further information about relationships between cells. An example of this is the Gabriel graph, a sub-graph of the Delaunay triangulation which can be generated following tessellation by connecting the cell centres with straight-line segments, which characterizes the proximity between cells (Fig. 1D). Each vertex of the Gabriel graph represents a cell, and the number of edges of each vertex gives the number of cell neighbours. The workflow of the image segmentation process is as follows: *Grouping Stage*: Image fragments of the same size, not necessarily disjoint, are grouped together based on similarity. This block-matching algorithm is less computationally demanding and is useful later on in the aggregation step. A fragment is grouped if its dissimilarity with a reference fragment falls below a specified threshold.
- *Collaborative Filtering*: Filtering is applied to each fragments group. A 3-dimensional linear transform is performed, followed by a transform-domain shrinkage (Wiener Filtering). The linear transform is then inverted to reproduce all (filtered) fragments.
- *Aggregation Stage*: The image is transformed back into its 2-dimensional form. All overlapping image fragments are weight-averaged to ensure that they are filtered for noise yet retain their distinct signal.

In this work we used BM3D software for image/video restoration and enhancement: Kostadin Dabov, Aram Danieyan, Alessandro Foi, Public release v2.00 (30 January 2014), Tampere University of Technology, http://www.cs.tut.fi/~foi/GCF-BM3D. Another, newer option for block-matching and 3D filtering is Targeted Image Denoising (TID): Enming Luo, TID - Targeted Image Denoising, MATLAB Central File Exchange. (29 August 2024), https://www.mathworks.com/matlabcentral/fileexchange/55776-tid-targeted-image-denoising.

1. Remove the image border and extract the meta-data from the image frame.
2. Read the grey-scale image from the file and use MATLAB function **im2double(imread(**‘I.jpg’)) to convert the image *I* to double precision and re-scale the output from an integer to be within the range [0,1]. Estimate the minimum, maximum, mean and standard deviation of the intensities.
3. Add white Gaussian noise at about 4% of the original image magnitude to smooth the image, before denoising the image. We use a block-matching and 3D filtering (BM3D) algorithm for noise reduction in images. An expansion of the non-local means methodology, there are two cascades in BM3D: a hard-thresholding stage and a Wiener filter stage, both involving the following parts: grouping, collaborative filtering, and aggregation.
  - *Grouping Stage*: Image fragments of the same size, not necessarily disjoint, are grouped together based on similarity. This block-matching algorithm is less computationally demanding and is useful later on in the aggregation step. A fragment is grouped if its dissimilarity with a reference fragment falls below a specified threshold.
  - *Collaborative Filtering*: Filtering is applied to each fragments group. A 3-dimensional linear transform is performed, followed by a transform-domain shrinkage (Wiener Filtering). The linear transform is then inverted to reproduce all (filtered) fragments.
  - *Aggregation Stage*: The image is transformed back into its 2-dimensional form. All overlapping image fragments are weight-averaged to ensure that they are filtered for noise yet retain their distinct signal.
  In this work we used BM3D software for image/video restoration and enhancement: Kostadin Dabov, Aram Danieyan, Alessandro Foi, Public release v2.00 (30 January 2014), Tampere University of Technology, http://www.cs.tut.fi/~foi/GCF-BM3D. Another, newer option for block-matching and 3D filtering is Targeted Image Denoising (TID): Enming Luo, TID - Targeted Image Denoising, MATLAB Central File Exchange. (29 August 2024), https://www.mathworks.com/matlabcentral/fileexchange/55776-tid-targeted-image-denoising.
4. For accurate detection of cell boundaries, we use an algorithm which is well known and widely used in medical applications for the detection of tube-like structures in an image, e.g., in the extraction of filamentous structures (blood vessels, nerves, neurites). It applies to both 2D and 3D images and was first described by Frangi et al. (28). The multi-scale second order local structure of an image (Hessian) is examined to develop a vessel enhancement filter, where a vesselness measure is obtained on the basis of all eigenvalues of the Hessian. This gives the excellent noise and background suppression and enhancement of filamentary structures in an image (29, 30).
5. To convert the gray-scale image to a binary image (a white background with back cell boundaries) the algorithm uses the MATLAB **imbinarize** function and a locally adaptive threshold, that is, choosing the threshold based on the local mean intensity in the neighborhood of each pixel. We have been able to select thresh-holding parameters that allow us to successfully process a wide range of corneal images at a depth 15—30μm automatically.
6. To fill tiny holes and create bridges when cell boundary pixels are close enough but not yet connected we use the MATLAB morphological close operation **imclose**. This performs an image dilation followed by erosion, with the same structuring element for both operations (a small disc of radius 1 pixel). The MATLAB function **bwareaopen** is then used to remove small artefacts from the binary image; in our case, we remove all connected objects with area less than 10 pixels, where the area was chosen experimentally.
7. As a final step we use a watershed transform, described on the MATLAB Image Processing Toolbox page, to segment an image. The watershed transform finds ‘catchment basins’ or ‘watershed ridge lines’ in an image by treating it as a surface, where light pixels represent high elevations and dark pixels represent low elevations. This method finds the centre of each object (using a morphological erode operation), calculates a distance map from the object centroids to their edges and then fills that ‘topological map’ with imaginary water, building a dam to separate the objects. The watershed algorithms in MATLAB and Fiji are identical, but image pre-processing in MATLAB, combined with the watershed approach, allows for high quality boundary identification when processing a large number of images automatically. In the future, we plan to apply machine learning to identify cell boundaries.
8. Our unique database allows us to separately consider the central and four peripheral regions of the cornea. Because the corners of the images are too dark to confidently identify cell boundaries and with the risk of segmentation errors affecting the numerical output, we include in the analysis only the cells that are completely in the central part of an image, i.e., a circle with a radius found experimentally, which corresponds to the brightest and most well-defined part of the image.
9. We then apply the MATLAB **regionprops** function to the binary image to measure properties such as area, centroid and bounding box, for each connected object (cell) in the image. The centroid is defined as the center of mass of the cell, returned as a 1-by-2 vector with horizontal (x-) coordinate and vertical (y-) coordinate denoting the center of mass.
10. A Gabriel Graph is then generated, based on the proximity of the cell centroids identified above (Fig. 1C).
11. All measured properties for each identified cell are written to a file.

A detailed description of the segmentation algorithm, its testing and comparison with alternatives will be published elsewhere. Statistical analysis of the output of the segmentation is carried out in RStudio.

### Uncertainties

Each of the probability distributions in Figs. 2 and 4 has a corresponding shaded region showing an estimate of the standard error of the distribution, calculated via the bootstrap method. The bootstrap standard error is calculated using the following steps (31):

1. Generate *N* samples (of length *n*, the number of cells in the image), with replacement, from the cell sample in an image.
2. For each of the *N* samples, calculate the probability distribution of the variable considered.
3. This results in *N* different estimates for the probability distribution of that variable. To find the bootstrapped standard error we take the standard error, *s*/√*n*, where *s* is the sample standard deviation, of these *N* probability estimates for each of the possible values of the variable.

The shaded regions in Fig. 3 are standard errors of the depth coefficient generated by the corresponding linear regression fit, carried out in RStudio.

## Statistical Testing

We employ the two-sample Kolmogorov--Smirnov test (32) to assess the similarity -- or otherwise -- of two probability distributions. As is standard we use the significance level *p=0*.*01*. When *p< 0*.*0*1, we reject the null hypothesis that the two samples have the same probability distribution (see Figs. **S4, S5, S8** and **S9** in **SI Appendix**), and conclude that the two populations are different. When *p>0*.*01* we accept the null hypothesis.

## Supporting information

SI Appendix for manuscript.

.RData and .R files for Figure Reproduction.

## Acknowledgments

We are grateful to Natalio Krasnogor and Yuchun Ding for useful discussions at an early stage of this work. We acknowledge the support of the following grants: MRC (G0900879) and Fight for Sight (5181/5182) and support from the Newcastle University UKRI Higher Education Innovation Fund (HEIF/H064).

