## Supplementary material for "Unlocking precision: How corneal cell area analysis revolutionizes post-transplant stem cell monitoring": SI Appendix for manuscript.

**This PDF file includes:**
Supporting text
Figs. S1 to S11
Tables S1 to S3

**Supporting Information Text**
The corneal epithelium is maintained by limbal stem cells (LSCs). Dysfunction of the LSCs, resulting from chemical and thermal burns, contact lens-related disease, congenial disorders, among other conditions, leads to limbal stem cell deficiency (LSCD), a sight-threatening condition. An effective treatment of LSCD, with 76% of patients reporting regained sight up to 24 months after the operation, consists of transplanting *ex-vivo* cultured LSCs from the patient's other healthy eye (i.e. autologous) or donor (i.e. allogeneic) to the affected eye. The post-operative assessment of corneal recovery is crucial but relies on ponderous and generally subjective visual inspection of a large number of microscopic images of the corneal epithelial cells, relying on the personal experience of the practitioner to interpret imprecise, qualitative diagnostic criteria. From a unique library of 100,000 cornea cell images from 34 patients, we have randomly selected 10 individuals (3,668 images) to demonstrate that the frequency distribution of the epithelial cell areas is a sensitive diagnostic tool of the corneal epithelium status. After a successful operation the distribution of cell areas is rather flat, reflecting an anomalously wide range of cell areas. As the cornea recovers, the frequency distribution becomes narrower with high statistical confidence and eventually approaches that of the healthy cornea. The corneal epithelial cell shape is independent of the cornea status despite a widespread expectation that healthy cells have a hexagonal shape. We also show that the corneal epithelial cell area distribution and its variation with the depth within the cornea are specific to each patient.

**Fig. S1.**

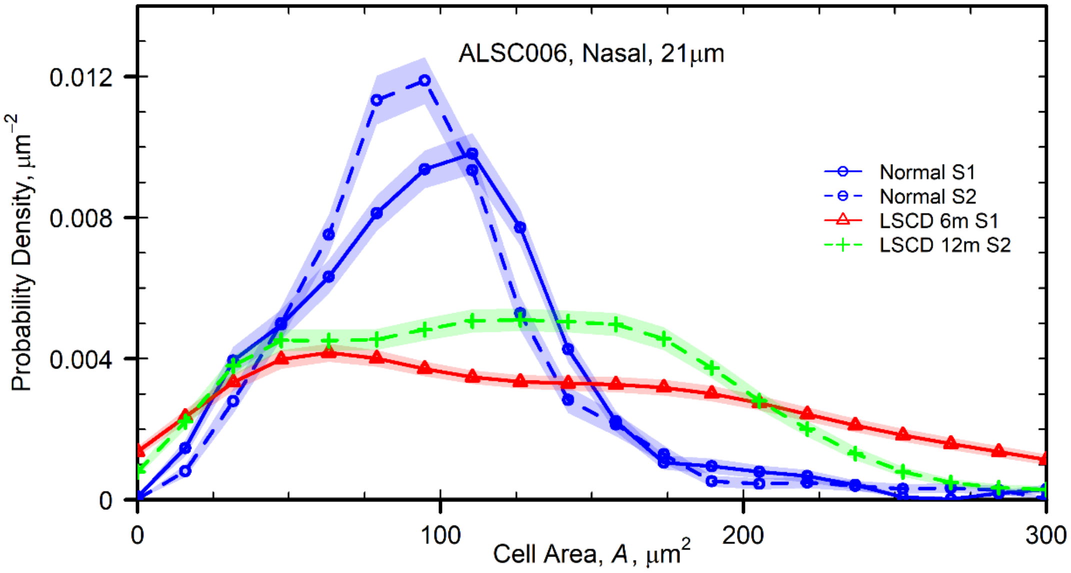


The PDF of cell areas of patient ALSC006 in the nasal corneal region at a depth of 21μm for the healthy (blue lines/circles) and post-operative central cornea at six months (red lines/triangles) and 12 months (green lines/plus signs) after surgery. The PDF for the normal eye has a pronounced maximum at 109μm$^2$ and a weak tail at large areas. The statistical difference of the PDFs at six, 12 and 24 months from the healthy distribution are assessed by the Kolmogorov-Smirnov test at the 1% significance level. The shaded regions show the bootstrap 2σ errors Materials and Methods for details). Labels S1--S2 indicate from what sequence of images these were taken, with PDFs differentiated by line type.

**Fig. S2.**

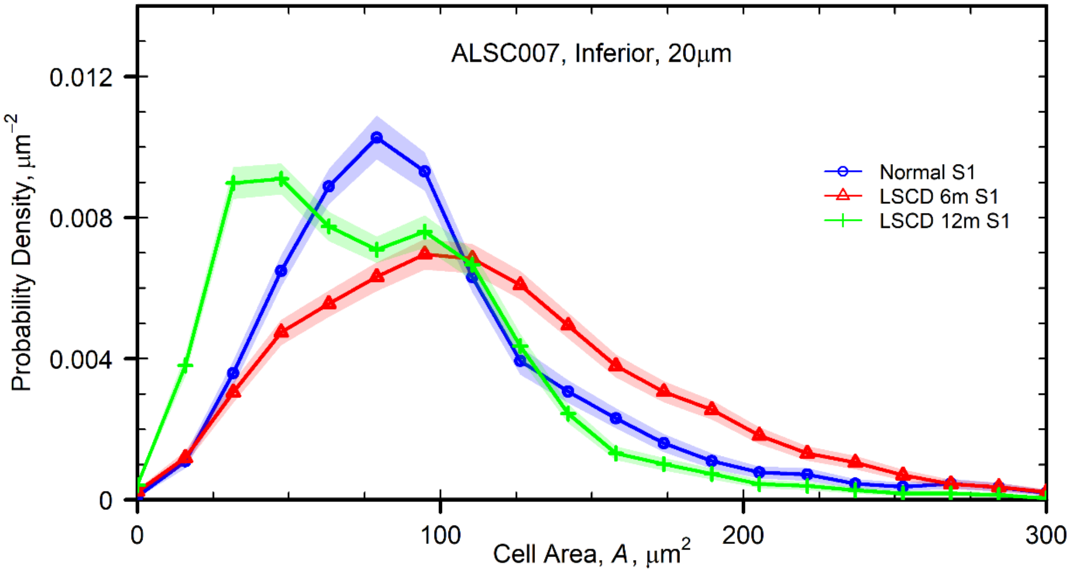


The PDF of cell areas of patient ALSC007 in the inferior corneal region at a depth of 20μM for the healthy (blue lines/circles) and post-operative central cornea at six months (red lines/triangles) and 12 months (green lines/plus signs) after surgery. The PDF for the normal eye has a pronounced maximum at 79$\mu m^2$ and a weak tail at large areas. The statistical difference of the PDFs at six and 12 months from the healthy distribution are assessed by the Kolmogorov-Smirnov test at the 1% significance level. The shaded regions show the bootstrap 2σ errors (see **Materials and Methods** for details).

**Figure S3.**

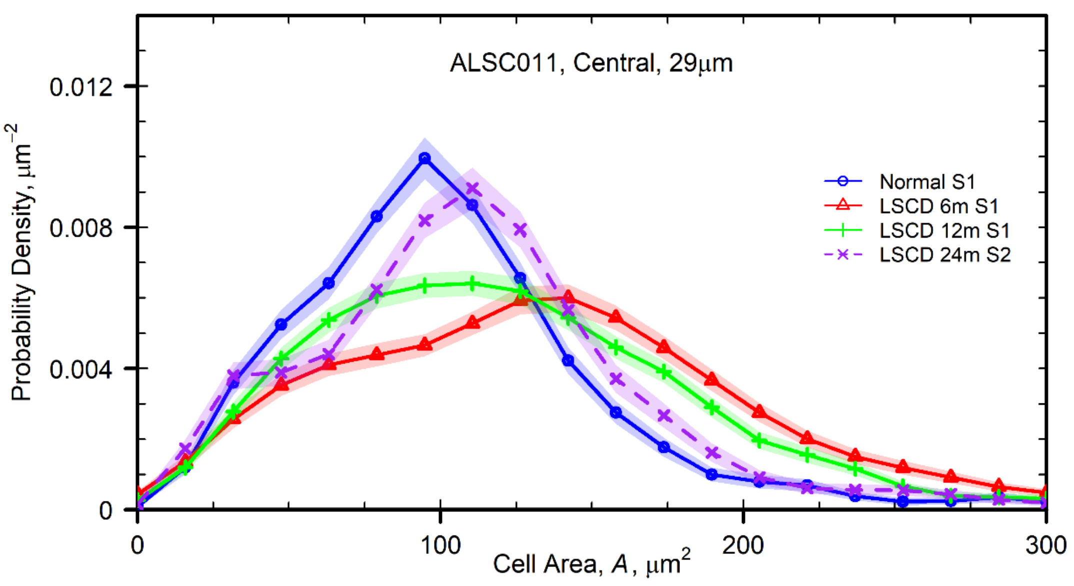


The PDF of cell areas of patient ALSC011 in the central corneal region at a depth of 29μm for the healthy (blue lines/circles) and post-operative central cornea at six months (red lines/triangles), 12 months (green lines/plus signs) and 24 months after surgery. The PDF for the normal eye has a pronounced maximum at 95μm$^2$ and a weak tail at large areas. The statistical difference of the PDFs at six, 12 and 24 months from the healthy distribution are assessed by the Kolmogorov-Smirnov test at the 1% significance level (see Fig. **S4**). The shaded regions show the bootstrap 2σ errors (see **Materials and Methods** for details). Labels S1--S2 indicate from what sequence of images these were taken, with PDFs differentiated by line type.

**Figure S4.**

Statistical Difference Testing: ALSC011, Central, 29um

| | Normal | LSCD6m | LSCD12m | LSCD24m |
|---|---|---|---|---|
| Normal | NA | Reject | Reject | Accept |
| LSCD6m | Reject | NA | Accept | Reject |
| LSCD12m | Reject | Accept | NA | Accept |
| LSCD24m | Accept | Reject | Accept | NA |

Table showing the results of Kolmogorov-Smirnov testing for distributions of cell areas in the central corneal epithelium of patient ALSC011 (Fig. **S3**). Testing was performed at 1% level of significance.

**Figure S5.**

Statistical Difference Testing: ALSC016, Nasal, 28um

| | Normal | LSCD6m | LSCD12m | LSCD24m |
|---|---|---|---|---|
| Normal | NA | Reject | Reject | Accept |
| LSCD6m | Reject | NA | Reject | Reject |
| LSCD12m | Reject | Reject | NA | Reject |
| LSCD24m | Accept | Reject | Reject | NA |

Table showing the results of Kolmogorov-Smirnov testing for distributions of cell areas in the central corneal epithelium of patient ALSC016 (Fig. **2A**). Testing was performed at 1% level of significance.

**Figure S6.**

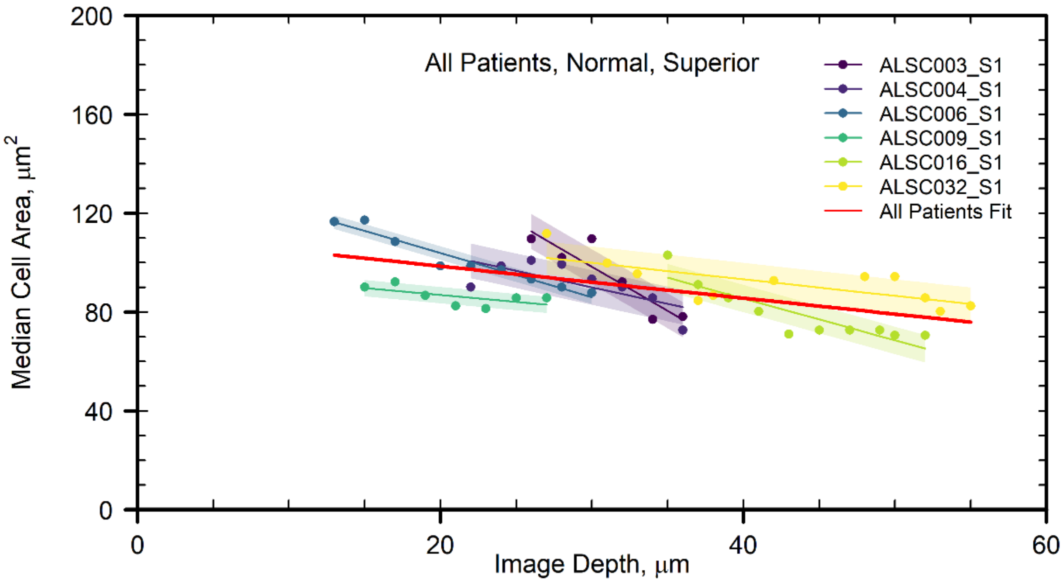


.

Distribution of median cell areas varying with depth for all patients in the healthy corneal epithelium, in the superior region. The coloured points are the individual medians at a given depth, for a given patient, with different patients distinguished by colour and different sequences distinguished by point shape. Red lines depict the linear relationship between depth and cell area, given by equation (1). This relationship was derived by performing a linear regression fit to all data points plotted. All other lines depict similar fits for each patient separately. The corresponding shaded regions show the 2σ uncertainty of the coefficient of depth for each fit. It is clear that all coefficients are negative. Depth is significant in the modelling of cell area in all fits at the 1% level (Table **S1**).

**Table S1.**

| Location | $A_0$ [μm²] | $\sigma_A$ [μm²] | $q$ [μm] | $\sigma_q$ [μm] | $p$ |
|---|---|---|---|---|---|
| Central | 129.1 | 8.6 | 1.0 | 0.3 | $9 \times 10^{-4}$ |
| Nasal | 132.1 | 3.8 | 1.2 | 0.1 | $9 \times 10^{-15}$ |
| Inferior | 143.5 | 7.8 | 1.6 | 0.3 | $1.4 \times 10^{-6}$ |
| Superior | 107.2 | 4.2 | 0.6 | 0.1 | $2 \times 10^{-6}$ |
| Temporal | 145.4 | 4.3 | 1.6 | 0.2 | $5.4 \times 10^{-16}$ |

The parameters of regression fit detailed in equation (1) fit to all patients in Figs. **3** and **S6** (red lines). The table shows, for each location, the values of $A_0$ and its standard deviation $\sigma_A$, the slope $q$ and its standard deviation $\sigma_q$ and the statistical significance $p$ of the fit.

**Table S2.**

| Patient/Sequence | $A_0$ [μm$^2$] | $\sigma_A$ [μm$^2$] | $q$ [μm] | $\sigma_q$ [μm] | $p$ |
|---|---|---|---|---|---|
| ALSC003/S1 | 217.3 | 22.9 | 3.1 | 0.6 | $3 \times 10^{-3}$ |
| ALSC005/S1 | 131.0 | 13.3 | 1.4 | 0.5 | $9 \times 10^{-2}$ |
| ALSC008/S1 | 169.7 | 9.5 | 2.6 | 0.5 | $1 \times 10^{-2}$ |
| ALSC008/S2 | 165.8 | 11.7 | 2.4 | 0.4 | $7 \times 10^{-5}$ |
| ALSC009/S1 | 84.6 | 3.2 | 0.4 | 0.1 | $9 \times 10^{-3}$ |
| ALSC011/S1 | 130.9 | 16.2 | 0.9 | 0.6 | 0.2 |
| ALSC016/S1 | 130.7 | 5.1 | 1.4 | 0.2 | $9 \times 10^{-5}$ |

The parameters of regression fit detailed in equation (1), fit to each individual patient in Fig. **3A**. The table shows, for each location, the values of $A_0$ and its standard deviation $\sigma_A$, the slope *q* and its standard deviation $\sigma_q$, and the statistical significance *p* of the fit.

### Figure S7.

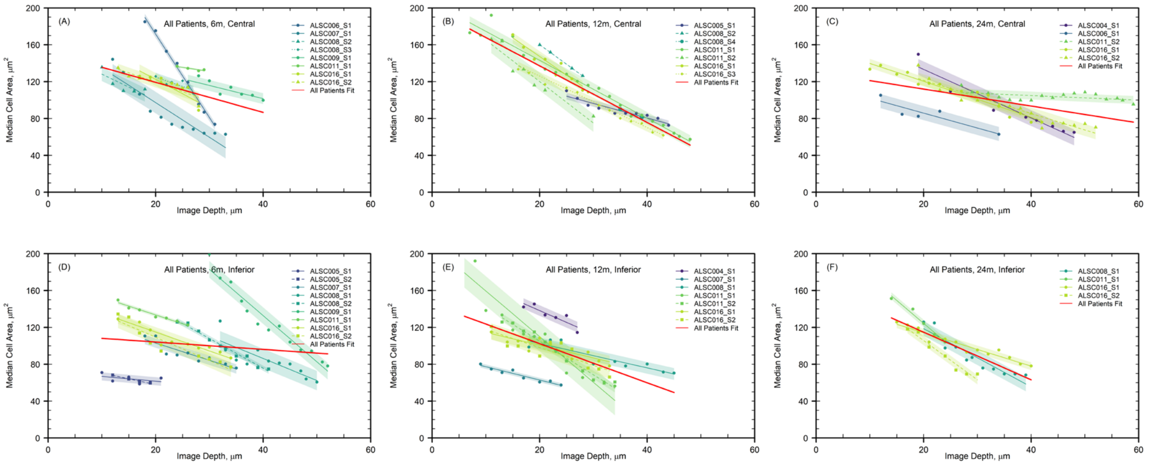


Distribution of median cell areas varying with depth for all patients in the central and inferior corneal epithelium, at (A,D) six, (B,E) 12 and (C,F) 24 months post-transplant. The coloured points are the individual medians at a given depth, for a given patient, with different patients distinguished by colour and different sequences distinguished by point shape. Red lines depict the linear relationship between depth and cell area, given by equation (1). This relationship was derived by performing a linear regression fit to all data points plotted. All other lines depict similar fits for each patient separately. The corresponding shaded regions show the 2σ uncertainty of the coefficient of depth for each fit. It is clear that all coefficients are negative. Depth is significant in the modelling of cell area in all fits at the 1% level (Table **S3**).

**Table S3.**

| Location/Condition | $A_0$ [μm²] | $\sigma_A$ [μm²] | $q$ [μm] | $\sigma_q$ [μm] | $p$ |
|---|---|---|---|---|---|
| Central/Normal | 129.1 | 8.6 | 1.0 | 0.3 | $9 \times 10^{-4}$ |
| Central/6m | 175.9 | 11.3 | 2.4 | 0.5 | $4 \times 10^{-6}$ |
| Central/12m | 198.6 | 4.1 | 3.1 | 0.1 | $2 \times 10^{-16}$ |
| Central/24m | 107.2 | 5.2 | 0.9 | 0.1 | $2 \times 10^{-2}$ |
| Inferior/Normal | 143.5 | 7.8 | 1.6 | 0.3 | $2 \times 10^{-6}$ |
| Inferior/6m | 112.2 | 9.0 | 0.4 | 0.3 | $2 \times 10^{-16}$ |
| Inferior/12m | 144.9 | 8.2 | 2.1 | 0.3 | $3 \times 10^{-8}$ |
| Inferior/24m | 166.5 | 7.6 | 1.6 | 0.3 | $9 \times 10^{-11}$ |

The parameters of regression fit detailed in equation (1) fit to all patients in Fig. **S7** (red lines). The table shows, for each location and condition, the values of $A_0$ and its standard deviation $\sigma_A$, the slope *q* and its standard deviation $\sigma_q$, and the statistical significance *p* of the fit. The entries labelled 'Normal' are identical to those in Table **S1**.

**Figure S8.**

Statistical Difference Testing: ALSC016, Central, 17um

| | Normal | LSCD6m_S1 | LSCD6m_S2 | LSCD12m | LSCD24m |
|---|---|---|---|---|---|
| Normal | NA | Accept | Accept | Accept | Accept |
| LSCD6m_S1 | Accept | NA | Accept | Accept | Accept |
| LSCD6m_S2 | Accept | Accept | NA | Accept | Accept |
| LSCD12m | Accept | Accept | Accept | NA | Accept |
| LSCD24m | Accept | Accept | Accept | Accept | NA |

Table showing the results of Kolmogorov-Smirnov testing for distributions of cell shapes in the corneal epithelium of patient ALSC016 (Fig. **4A**). Testing was performed at 1% level of significance.

**Figure S9.**

Statistical Difference Testing: ALSC011, Central, 29um

| | Normal | LSCD6m | LSCD12m | LSCD24m |
|---|---|---|---|---|
| Normal | NA | Accept | Accept | Accept |
| LSCD6m | Accept | NA | Accept | Accept |
| LSCD12m | Accept | Accept | NA | Accept |
| LSCD24m | Accept | Accept | Accept | NA |

Table showing the results of Kolmogorov-Smirnov testing for distributions of cell shapes in the corneal epithelium of patient ALSC011 (Fig. **4B**). Testing was performed at 1% level of significance.

**Figure S10.**

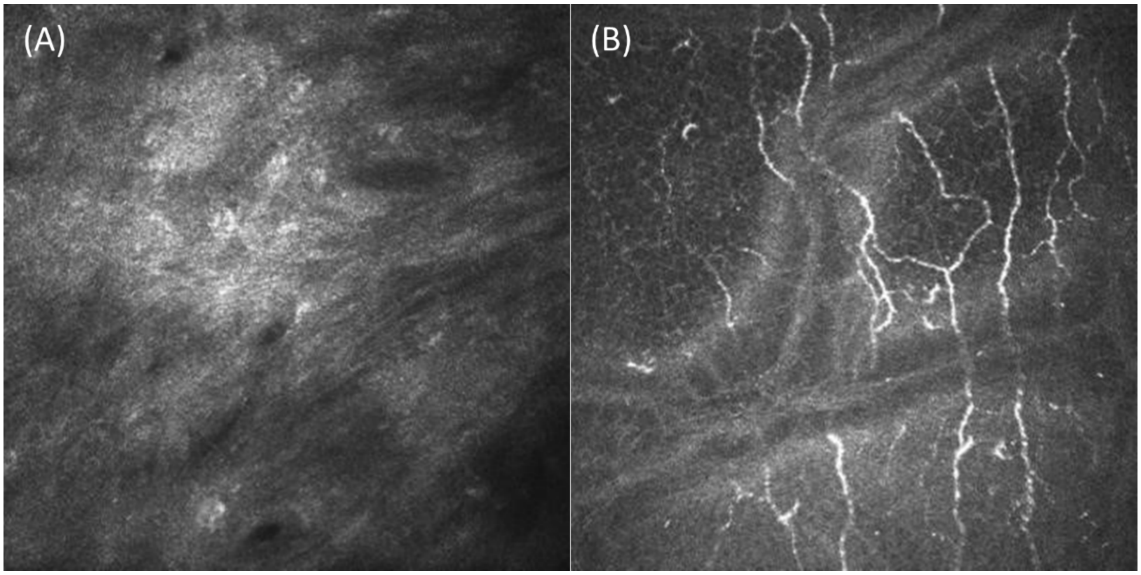


(A) IVCM image of the LSCD affected cornea pre-treatment (Baseline). Significant lesioning of the corneal epithelium leaves little to no identifiable cell structure, making these images unsuitable for morhological analysis. (B) IVCM image of the normal (healthy cornea) at a depth of 64μm. At this depth, beyond the corneal epithelium, sub-epithelial nerves can be seen.

**Figure S11.**

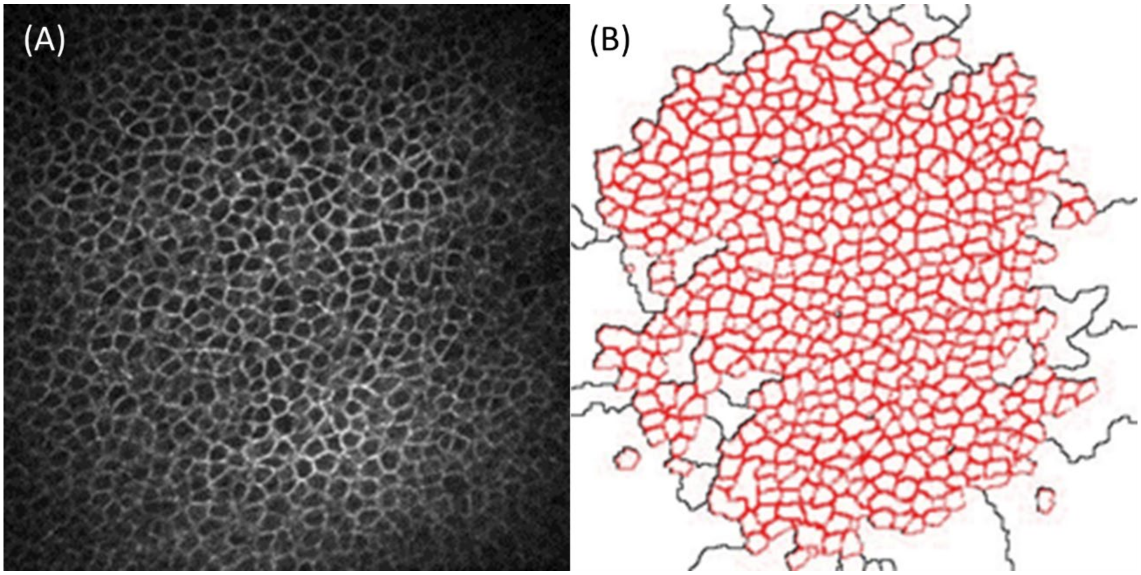


Identification of cell boundaries using the ImageJ segmentation algorithm. (A) Original IVCM image of central corneal epithelium. (B) Identification of cell boundaries.
