## Supplementary material for "Unlocking precision: How corneal cell area analysis revolutionizes post-transplant stem cell monitoring": .RData and .R files for Figure Reproduction.: Figure 2A.R

```
#Figure 2A.
#Load 'Figure 2A.RData' before running.
#Relevant Functions:
#Function for generating bootstrap sample of PDFs.
bootstrap = function(x){
  bootSample = replicate(1000, sample(eval(as.name(x)), replace=T))
  bootSample1 = numeric(1000)
  bootSampleMean = numeric(1000)
  bootSD = numeric(20)
  bootMean = numeric(20)
  for (i in 1:20) {
    for (j in 1:1000) {
      bootSampleMean[j] = mean(bootSample[,j])
      bootsampleDensity = density(bootSample[,j], n=20, from = 0, to = 300)$y
      bootSample1[j] =  bootsampleDensity[i]
    }
    bootSD[i] = sd(bootSample1)
    bootMean[i] = mean(bootSample1)
  }
  bootMeanSE = sd(bootSampleMean)
  return(list(bootSD, bootMean, bootMeanSE))
}
#Function for carrying out Mann-Whitney and Kolmogorov-Smirnoff Tests.
#Accept meaning 'Accept the Null Hypothesis, PDFs of subsequent depths are statistically similar at the 5% level.'
#Reject meaning 'Reject the Null Hypothesis, PDFs of subsequent depths are NOT statistically similar at the 5% level.'
StatisticalDifferenceTest = function(WorkingDF, WorkingDFPlus1){
  #Mann-Whitney & Kolmogorov Smirnoff Testing.
  MWTest = wilcox.test(WorkingDF, WorkingDFPlus1, alternative = "two.sided")
  MW_p = MWTest$p.value
  #Kolmogorov Smirnoff Testing!!!
  KSTest = ks.test(WorkingDF, WorkingDFPlus1, alternative = "two.sided")
  KS_p = KSTest$p.value
  if(MW_p > 0.05 & KS_p > 0.05){
    Output = 'Accept'
  }else if(MW_p > 0.05 & KS_p <= 0.05) {
    Output = 'KS Reject'
  }else if(MW_p <= 0.05 & KS_p > 0.05) {
    Output = 'MW Reject'
  }else{
    Output = 'Reject'
  }
  return(Output)
}
#PDFs generated using density() function with n = 20.
#Setting colours for plotting. 
colours = c('blue', 'red', 'green', 'purple')
#Setting plot variables (Patient No., Corneal Location, Depth (microns))
patient = 'ALSC016'
location = 'Central'
depth = 25
#Now we can begin plotting:
#Plot Cell Area axis ranges from 0:300, plotting at intervals of 25.
#Probability Density axis range set as 0:0.12 at intervals of 0.01.
plot(x = seq(0, 300, 25), y = seq(0,0.12, 0.01), col = 'white',type = 'l', lty = 1, lwd = 1, 
     main = '', 
     ylab = '', 
     xlab = '',
     ylim = c(0,0.014), yaxs= 'i', xaxs = 'i', xaxt = 'n', yaxt = 'n',
     family = "font")
#Titleing plot using variables set above.
title(main = bquote(paste( .(patient),", ", .(location), ", ", .(depth), mu, "m", sep = '')), line = -2)
#Axes labels.
title(ylab = expression(paste("Probability Density, ",italic(mu), m^-2)), line =4)
title(xlab = expression(paste("Cell Area, ", italic(A), ", ", italic(mu), m^2)), line = 2)
#Axes labels and tickmarks added.
axis(side = 1, at = c(0,100,200,300,400,500), labels = c(0,100,200,300,400,500))
rug(x = seq(0,300, 100), ticksize = 0.02)
rug(x = seq(0,300, 100/4), ticksize = 0.015)
axis(side = 2, at = c(0,0.004 , 0.008, 0.012, 0.016, 0.02), labels = c(0,0.004 , 0.008, 0.012, 0.016, 0.02), las = 2)
rug(x = seq(0,0.02, 0.004/4), ticksize = 0.015, side = 2)
rug(x = seq(0,0.02, 0.004), ticksize = 0.02, side = 2)
rug(x = seq(0,300, 100/4), ticksize = 0.015, side = 3)
rug(x = seq(0,0.02, 0.004/4), ticksize = 0.015, side = 4)
#Initialising variables to store details of PDFs plotted for legend.
PatientList_Sequence = vector()
ColoursVector = vector()
SymbolVector = vector()
LineTypeVector = vector()
#We plot each available condition for the given patient and location. 
#Initialising all possible conditions.
ConditionList = c('Normal', '6m', '12m', '24m')
#We plot each available PDF for the given condition. These are differentiated by
#a 'Sequence No.'.
#Initialising all possible sequences.
SequenceList = c('S1', 'S2', 'S3', 'S4', 'S5', 'S6')
#Plotting PDF of given Normal (healthy) and LSCD affected cornea at 6, 12 and 24
#months post-treatment (with bootstrap error).
for (k in 1:length(ConditionList)){
  #Set condition to plot.
  condition = ConditionList[k]
for (l in 1:length(SequenceList)){
  #Set sequence number.
  sequence = SequenceList[l]
  #Begin by plotting the bootstrap error. 
  #Check if that sequence number exists for the given patient, location and condition. 
  if(exists(paste(patient,'_', location, '_', condition, '_', sequence, '_pdf_', depth, sep =''))){
    #Assign the corresponding PDF variable to a placeholder 'WorkingDF'.
    WorkingDF = eval(as.name(paste(patient,'_', location, '_', condition, '_', sequence, '_pdf_', depth, sep ='')))
    #Assign the corresponding bootstrap error ('boot') variable to a placeholder 'WorkingBoot'.
    WorkingBoot = eval(as.name(paste(patient,'_', location, '_', condition, '_', sequence, '_boot_', depth, sep ='')))
    #Plot the error bar (a shaded region), for the PDF. 
    polygon(x = c(WorkingDF$x, rev(WorkingDF$x)),
            y = c(WorkingDF$y - WorkingBoot[[1]], 
                  rev(WorkingDF$y + WorkingBoot[[1]])),
            col = adjustcolor(colours[k], alpha.f=0.2), border = NA)
  }
  #Now plotting the PDF.
  #Again check if that sequence number exists for the given patient, location and condition. 
  if(exists(paste(patient,'_', location, '_', condition, '_', sequence, '_pdf_', depth, sep =''))){
    #Storing line name of this PDF for Legend.
    PatientList_Sequence = c(PatientList_Sequence, paste(condition, ' ', sequence, sep =''))
    #Plot the PDF, with colour and line marker corresponding to condition plotting. Line Type corresponds to sequence number. 
    lines(eval(as.name(paste(patient,'_', location, '_', condition, '_', sequence, '_pdf_', depth, sep =''))), type = 'o', 
          col = colours[k], lty = l, lwd = 1, pch = k)
    #Storing colour of this PDF for Legend.
    ColoursVector = c(ColoursVector, colours[(k)])
    #Storing line marker of this PDF for Legend.
    SymbolVector = c(SymbolVector, k)
    #Storing line type of this PDF for Legend.
    LineTypeVector = c(LineTypeVector, l)
  }
}
  #Repeat for all available conditions.
}
if(length(PatientList_Sequence)!=0){
  legend(x = 225, y = 0.01, 
         legend = PatientList_Sequence, 
         col = ColoursVector,
         pch = SymbolVector,
         lwd = rep(0.5, length(PatientList_Sequence)),
         lty = LineTypeVector,        
         bty="n", cex = 1)
}
```
