## Supplementary material for "Unlocking precision: How corneal cell area analysis revolutionizes post-transplant stem cell monitoring": .RData and .R files for Figure Reproduction.: Figure 3A.R

```
#Figure 3A.
#Load 'Figure 3A.RData' before running.
library("plyr")
library("viridis")
#Setting colours for plotting. 
colours = viridis(10)
#Setting plot variables (Condition & Location).
Condition = 'Normal'
Location = 'Central'
#Now we can begin plotting:
#Plot Image Depth axis ranges from 0:60, plotting at each available depth.
#Median Cell Area axis range set as 0:200.
plot(x = c(1,100), y = c(1,100), col = 'white',type = 'l', lty = 1, lwd = 1, 
         main = '', 
         ylab = '', 
         xlab = '',
         xlim = c(0,60),
         ylim = c(0,200), yaxs= 'i', xaxs = 'i', yaxt = 'n', xaxt = 'n',
         family = "font")
#Titleing plot using variables set above.
title(main = bquote(paste( "All Patients, ", .(Condition), ", ",.(Location), sep = '')), line = -2) 
#Axes labels.
title(ylab = expression(paste("Median Cell Area, ",italic(mu), m^2)), line =4)
title(xlab = expression(paste("Image Depth, ", italic(mu), m)), line = 2)
#Axes labels and tickmarks added.
axis(side = 1, at = seq(round_any(min(strtoi(WorkingImageDepths)), 20, f = floor),round_any(max(strtoi(WorkingImageDepths)), 20, f = ceiling),20), labels = seq(round_any(min(strtoi(WorkingImageDepths)), 20, f = floor),round_any(max(strtoi(WorkingImageDepths)), 20, f = ceiling),20))
rug(x = seq(0, 60, 20/4), ticksize = 0.02)
axis(side = 2, at = seq(0,400,40),labels = seq(0,400,40), las = 2)
rug(x = seq(0,200,40/4), ticksize = 0.015, side = 2)
rug(x = seq(0,200,40), ticksize = 0.02, side = 2)
rug(x = seq(0, 60, 20/4), ticksize = 0.015, side = 3)
rug(x = seq(0,200,40/4), ticksize = 0.015, side = 4)
#Once plot set, we add Mean Error bars (if present).
#Plotting error bars as +/- Standard Error of PDF of Mean. 
#Initialising variables to store details of PDFs plotted for legend.
PatientList_Sequence = vector()
ColoursVector = vector()
ColoursVectorPatients = vector()
SymbolVector = vector()
LineTypeVector = vector()
#We plot median cell area at all depths for each available patient sequence for the given condition and location. 
for(k in 1:length(PatientList)){
  for (l in 1:length(SequenceList)) {
    #Set sequence number.
    Sequence = SequenceList[l]
    if(exists(paste(PatientList[k],'_', Location, '_', Condition, '_', Sequence, '_median', sep =''))){
      #Begin by plotting error of the linear regression fit. 
      #Check if that sequence number exists for the given patient, location and condition. 
      WorkingDF = eval(as.name(paste(PatientList[k],'_', Location, '_', Condition, '_', Sequence, '_median', sep ='')))
      WorkingDF$ImageDepthCon = as.numeric(WorkingDF$ImageDepthCon)
      #Fit the linear relationship between cell area median and image depth.
      model = lm(WorkingMedian ~ ImageDepthCon, data = WorkingDF)
      #Assign the corresponding fit error to a placeholder 'WorkingError'.
      WorkingError = summary(model)$sigma
      x = WorkingDF$ImageDepthCon
      y = model$coefficients[2]*x + model$coefficients[1]
      #Plot the error bar (a shaded region), for the linear regression fit. 
      polygon(x = c(x, rev(x)),
              y = c(y - WorkingError, 
                    rev(y + WorkingError)),
              col = adjustcolor(colours[(k)], alpha.f=0.2), border = NA)
    }
        
  }
}
#Now plotting the point medians and linear relationship.
#Again check if that sequence number exists for the given patient, location and condition. 
for (k in  1:length(PatientList)) {
  for (l in 1:length(SequenceList)) {
    #Set sequence number.
    Sequence = SequenceList[l]
    #Again check if that sequence number exists for the given patient, location and condition. 
    if(exists(paste(PatientList[k], '_', Location, '_', Condition, '_', Sequence, '_median', sep =''))){
      #Assign the corresponding median variable to a placeholder 'WorkingDF'.
      WorkingDF = eval(as.name(paste(PatientList[k], '_', Location, '_', Condition, '_', Sequence, '_median', sep ='')))
      WorkingDF = WorkingDF[order(strtoi(WorkingDF$ImageDepthCon)),]
      WorkingDF$ImageDepthCon = as.numeric(WorkingDF$ImageDepthCon)
      #Fit the linear relationship between cell area median and image depth.
      model = lm(WorkingMedian ~ ImageDepthCon, data = WorkingDF)
      #Plot the individual cell area medians as points. 
      points(x = WorkingDF$ImageDepthCon, y = WorkingDF$WorkingMedian, 
             col = colours[(k)], bg = colours[(k)], pch = (l%%length(SequenceList)+15))
      #Plot the linear relationship from 'model'.
      lines(x = WorkingDF$ImageDepthCon, y = WorkingDF$ImageDepthCon*model$coefficients[2]+model$coefficients[1], col = colours[(k)], lty = l, lwd = 0.5)
      #Storing colour of these plotted medians for Legend.
      ColoursVector = c(ColoursVector, colours[(k)])
      #Storing markers of these plotted medians for Legend.
      SymbolVector = c(SymbolVector, (l%%length(SequenceList)+15))
      #Storing line type of the linear relationship for Legend.
      LineTypeVector = c(LineTypeVector, l)
      #Storing names of these points and lines for Legend.
      PatientList_Sequence = c(PatientList_Sequence, paste(PatientList[k],'_', Sequence,": a = ", signif(model$coefficients[2], 3), ", b = ", signif(model$coefficients[1], 3), ", sd = ", signif(summary(model)$sigma, 3), " p =", signif(summary(model)$coefficients[8], 3), sep =''))
      }
    }
}
#Now we perform the ame linear regression fit to all cell area medians plotted, 
#regardless of patient.
#Initialise variable to store all medians plotted.
AllMedians = data.frame()
#Function to populate 'AllMedians' with every plotted cell area median.
for (k in  1:length(PatientList)) {
  for (l in 1:length(SequenceList)) {
    Sequence = SequenceList[l]
    if(exists(paste(PatientList[k], '_', Location, '_', Condition, '_', Sequence, '_median', sep =''))){
      WorkingDF = eval(as.name(paste(PatientList[k], '_', Location, '_', Condition, '_', Sequence, '_median', sep ='')))
      WorkingDF = WorkingDF[order(strtoi(WorkingDF$ImageDepthCon)),]
      AllMedians = rbind(AllMedians, WorkingDF)
    }
  }
}
#Ordering ALL Medians.
AllMedians = AllMedians[order(strtoi(AllMedians$ImageDepthCon)),]
AllMedians$ImageDepthCon = strtoi(AllMedians$ImageDepthCon)
#Fit linear relationship between ALL Medians. 
model5 = lm(formula = WorkingMedian ~ ImageDepthCon, data = AllMedians)
lines(AllMedians$ImageDepthCon, predict(model5), col= 'red', lwd = 1)
if(length(PatientList_Sequence!=0)){
  legend(x = 45, y = 190, 
          legend = c(substring(PatientList_Sequence, 1, 10), 'All Patients Fit'), 
          col = c(ColoursVector, 'red'),
          bg = c(ColoursVector, 'red'),
          pch = c(SymbolVector, NA),
          lwd = rep(0.5, length(PatientList_Sequence)),
          lty = c(LineTypeVector, 1),         
          bty="n", cex = 1)
}
```
