## Supplementary material for "Unlocking precision: How corneal cell area analysis revolutionizes post-transplant stem cell monitoring": .RData and .R files for Figure Reproduction.: README.txt

READ ME

========================
General Information
========================

Author: Patrick Parkinson
Contact:
DOI: https://doi.org/10.25405/data.ncl.26954446
License:
Last updated:23/08/2024
Related article: Unlocking Precision: How Corneal Cell Area Analysis Revolutionizes Post-Transplant Stem Cell Monitoring

========================
Introductory information
========================
Files included in the data deposit (include a short description of what data are contained): 

1) Figure 2A.R - R Script for reproducing Figure 2A from 'Unlocking Precision: How Corneal Cell Area Analysis Revolutionizes Post-Transplant Stem Cell Monitoring'.
2) Figure 2A.RData - R Data required for reproducing Figure 2A from 'Unlocking Precision: How Corneal Cell Area Analysis Revolutionizes Post-Transplant Stem Cell Monitoring'.
3) Figure 2B.R - R Script for reproducing Figure 2B from 'Unlocking Precision: How Corneal Cell Area Analysis Revolutionizes Post-Transplant Stem Cell Monitoring'.
4) Figure 2B.RData - R Data for reproducing Figure 2B from 'Unlocking Precision: How Corneal Cell Area Analysis Revolutionizes Post-Transplant Stem Cell Monitoring'.
5) Figure 2C.R - R Script for reproducing Figure 2C from 'Unlocking Precision: How Corneal Cell Area Analysis Revolutionizes Post-Transplant Stem Cell Monitoring'.
6) Figure 2C.RData - R Data for reproducing Figure 2C from 'Unlocking Precision: How Corneal Cell Area Analysis Revolutionizes Post-Transplant Stem Cell Monitoring'.
7) Figure 2D.R - R Script for reproducing Figure 2D from 'Unlocking Precision: How Corneal Cell Area Analysis Revolutionizes Post-Transplant Stem Cell Monitoring'.
8) Figure 2D.RData - R Data for reproducing Figure 2D from 'Unlocking Precision: How Corneal Cell Area Analysis Revolutionizes Post-Transplant Stem Cell Monitoring'.
9) Figure 3A.R - R Script for reproducing Figure 3A from 'Unlocking Precision: How Corneal Cell Area Analysis Revolutionizes Post-Transplant Stem Cell Monitoring'.
10) Figure 3A.RData - R Data for reproducing Figure 3A from 'Unlocking Precision: How Corneal Cell Area Analysis Revolutionizes Post-Transplant Stem Cell Monitoring'.
11) Figure 3B.R - R Script for reproducing Figure 3B from 'Unlocking Precision: How Corneal Cell Area Analysis Revolutionizes Post-Transplant Stem Cell Monitoring'.
12) Figure 3B.RData - R Data for reproducing Figure 3B from 'Unlocking Precision: How Corneal Cell Area Analysis Revolutionizes Post-Transplant Stem Cell Monitoring'.
13) Figure 3C.R - R Script for reproducing Figure 3C from 'Unlocking Precision: How Corneal Cell Area Analysis Revolutionizes Post-Transplant Stem Cell Monitoring'.
14) Figure 3C.RData - R Data for reproducing Figure 3C from 'Unlocking Precision: How Corneal Cell Area Analysis Revolutionizes Post-Transplant Stem Cell Monitoring'.
15) Figure 3D.R - R Script for reproducing Figure 3D from 'Unlocking Precision: How Corneal Cell Area Analysis Revolutionizes Post-Transplant Stem Cell Monitoring'.
16) Figure 3D.RData - R Data for reproducing Figure 3D from 'Unlocking Precision: How Corneal Cell Area Analysis Revolutionizes Post-Transplant Stem Cell Monitoring'.
17) Figure 4A.R - R Script for reproducing Figure 4A from 'Unlocking Precision: How Corneal Cell Area Analysis Revolutionizes Post-Transplant Stem Cell Monitoring'.
18) Figure 4A.RData - R Data for reproducing Figure 4A from 'Unlocking Precision: How Corneal Cell Area Analysis Revolutionizes Post-Transplant Stem Cell Monitoring'.
19) Figure 4B.R - R Script for reproducing Figure 4B from 'Unlocking Precision: How Corneal Cell Area Analysis Revolutionizes Post-Transplant Stem Cell Monitoring'.
20) Figure 4B.RData - R Data for reproducing Figure 4B from 'Unlocking Precision: How Corneal Cell Area Analysis Revolutionizes Post-Transplant Stem Cell Monitoring'.
21) Figure 4C.R - R Script for reproducing Figure 4C from 'Unlocking Precision: How Corneal Cell Area Analysis Revolutionizes Post-Transplant Stem Cell Monitoring'.
22) Figure 4C.RData - R Data for reproducing Figure 4C from 'Unlocking Precision: How Corneal Cell Area Analysis Revolutionizes Post-Transplant Stem Cell Monitoring'.
23) Figure 4D.R - R Script for reproducing Figure 4D from 'Unlocking Precision: How Corneal Cell Area Analysis Revolutionizes Post-Transplant Stem Cell Monitoring'.
24) Figure 4D.RData - R Data for reproducing Figure 4D from 'Unlocking Precision: How Corneal Cell Area Analysis Revolutionizes Post-Transplant Stem Cell Monitoring'.
25) Figure S1.R - R Script for reproducing Figure S1 from 'Unlocking Precision: How Corneal Cell Area Analysis Revolutionizes Post-Transplant Stem Cell Monitoring - SI Appendix'.
26) Figure S1.RData - R Data for reproducing Figure S1 from 'Unlocking Precision: How Corneal Cell Area Analysis Revolutionizes Post-Transplant Stem Cell Monitoring - SI Appendix'.
27) Figure S2.R - R Script for reproducing Figure S2 from 'Unlocking Precision: How Corneal Cell Area Analysis Revolutionizes Post-Transplant Stem Cell Monitoring - SI Appendix'.
28) Figure S2.RData - R Data for reproducing Figure S2 from 'Unlocking Precision: How Corneal Cell Area Analysis Revolutionizes Post-Transplant Stem Cell Monitoring - SI Appendix'.
29) Figure S3.R - R Script for reproducing Figure S3 from 'Unlocking Precision: How Corneal Cell Area Analysis Revolutionizes Post-Transplant Stem Cell Monitoring - SI Appendix'.
30) Figure S3.RData - R Data for reproducing Figure S3 from 'Unlocking Precision: How Corneal Cell Area Analysis Revolutionizes Post-Transplant Stem Cell Monitoring - SI Appendix'.
31) Figure S4.R - R Script for reproducing Figure S4 from 'Unlocking Precision: How Corneal Cell Area Analysis Revolutionizes Post-Transplant Stem Cell Monitoring - SI Appendix'.
32) Figure S4.RData - R Data for reproducing Figure S4 from 'Unlocking Precision: How Corneal Cell Area Analysis Revolutionizes Post-Transplant Stem Cell Monitoring - SI Appendix'.
33) Figure S5.R - R Script for reproducing Figure S5 from 'Unlocking Precision: How Corneal Cell Area Analysis Revolutionizes Post-Transplant Stem Cell Monitoring - SI Appendix'.
34) Figure S5.RData - R Data for reproducing Figure S5 from 'Unlocking Precision: How Corneal Cell Area Analysis Revolutionizes Post-Transplant Stem Cell Monitoring - SI Appendix'.
35) Figure S6.R - R Script for reproducing Figure S6 from 'Unlocking Precision: How Corneal Cell Area Analysis Revolutionizes Post-Transplant Stem Cell Monitoring - SI Appendix'.
36) Figure S6.RData - R Data for reproducing Figure S6 from 'Unlocking Precision: How Corneal Cell Area Analysis Revolutionizes Post-Transplant Stem Cell Monitoring - SI Appendix'.
37) Figure S7(A).R - R Script for reproducing Figure S7(A) from 'Unlocking Precision: How Corneal Cell Area Analysis Revolutionizes Post-Transplant Stem Cell Monitoring - SI Appendix'.
38) Figure S7(A).RData - R Data for reproducing Figure S7(A) from 'Unlocking Precision: How Corneal Cell Area Analysis Revolutionizes Post-Transplant Stem Cell Monitoring - SI Appendix'.
39) Figure S7(B).R - R Script for reproducing Figure S7(B) from 'Unlocking Precision: How Corneal Cell Area Analysis Revolutionizes Post-Transplant Stem Cell Monitoring - SI Appendix'.
40) Figure S7(B).RData - R Data for reproducing Figure S7(B) from 'Unlocking Precision: How Corneal Cell Area Analysis Revolutionizes Post-Transplant Stem Cell Monitoring - SI Appendix'.
41) Figure S7(C).R - R Script for reproducing Figure S7(C) from 'Unlocking Precision: How Corneal Cell Area Analysis Revolutionizes Post-Transplant Stem Cell Monitoring - SI Appendix'.
42) Figure S7(C).RData - R Data for reproducing Figure S7(C) from 'Unlocking Precision: How Corneal Cell Area Analysis Revolutionizes Post-Transplant Stem Cell Monitoring - SI Appendix'.
43) Figure S7(D).R - R Script for reproducing Figure S7(D) from 'Unlocking Precision: How Corneal Cell Area Analysis Revolutionizes Post-Transplant Stem Cell Monitoring - SI Appendix'.
44) Figure S7(D).RData - R Data for reproducing Figure S7(D) from 'Unlocking Precision: How Corneal Cell Area Analysis Revolutionizes Post-Transplant Stem Cell Monitoring - SI Appendix'.
45) Figure S7(E).R - R Script for reproducing Figure S7(E) from 'Unlocking Precision: How Corneal Cell Area Analysis Revolutionizes Post-Transplant Stem Cell Monitoring - SI Appendix'.
46) Figure S7(E).RData - R Data for reproducing Figure S7(E) from 'Unlocking Precision: How Corneal Cell Area Analysis Revolutionizes Post-Transplant Stem Cell Monitoring - SI Appendix'.
47) Figure S7(F).R - R Script for reproducing Figure S7(F) from 'Unlocking Precision: How Corneal Cell Area Analysis Revolutionizes Post-Transplant Stem Cell Monitoring - SI Appendix'.
48) Figure S7(F).RData - R Data for reproducing Figure S7(F) from 'Unlocking Precision: How Corneal Cell Area Analysis Revolutionizes Post-Transplant Stem Cell Monitoring - SI Appendix'.
49) Figure S8.R - R Script for reproducing Figure S8 from 'Unlocking Precision: How Corneal Cell Area Analysis Revolutionizes Post-Transplant Stem Cell Monitoring - SI Appendix'.
50) Figure S8.RData - R Data for reproducing Figure S8 from 'Unlocking Precision: How Corneal Cell Area Analysis Revolutionizes Post-Transplant Stem Cell Monitoring - SI Appendix'.
49) Figure S9.R - R Script for reproducing Figure S9 from 'Unlocking Precision: How Corneal Cell Area Analysis Revolutionizes Post-Transplant Stem Cell Monitoring - SI Appendix'.
50) Figure S9.RData - R Data for reproducing Figure S9 from 'Unlocking Precision: How Corneal Cell Area Analysis Revolutionizes Post-Transplant Stem Cell Monitoring - SI Appendix'.

Explain the relationship between multiple data sets, if required:

========================== 
Methodological information
==========================
A brief method description � what the data is, how and why it was collected or created, and how it was processed:

All '.RData' files contain data extracted from in-vivo confocal microscopy (IVCM) images of the human corneal epithelium by means of an image analysis algorithm developed in MATLAB.
For each cell in a given image, the algorithm extracts it's area and number of neighbours (both relevant for this publication), as well as a number of other metrics.
The output of this image analysis comes in the form of a .txt file, listing these details for each cell.
The output is then read into a '.RData' file, before being analysed in RStudio.

Instruments, hardware and software used: 
In-Vivo Confocal Microscope (Operated by Francisco Figueredo)
MATLAB
RStudio

Date(s) of data collection: 


Geographic coverage of data:
N/A


Data validation (how was the data checked, proofed and cleaned):


Overview of secondary data, if used:


=========================
Data-specific information
=========================
Definitions of names, labels, acronyms or specialist terminology uses for variables, records and their values: 


Explanation of weighting and grossing variables: 


Outline any missing data:
