## Supplementary material for "Unlocking precision: How corneal cell area analysis revolutionizes post-transplant stem cell monitoring": .RData and .R files for Figure Reproduction.: Figure S4.R

```
#Figure S4.
#Load 'Figure S4.RData' before running.
StatisticalDifferenceTest = function(WorkingDF, WorkingDFPlus1){
  #Mann-Whitney & Kolmogorov Smirnoff Testing.
  MWTest = wilcox.test(WorkingDF, WorkingDFPlus1, alternative = "two.sided")
  MW_p = MWTest$p.value
  #Kolmogorov Smirnoff Testing!!!
  KSTest = ks.test(WorkingDF, WorkingDFPlus1, alternative = "two.sided")
  KS_p = KSTest$p.value
  if(MW_p > 0.0001 & KS_p > 0.0001){
    Output = 'Accept'
  }else if(MW_p > 0.0001 & KS_p <= 0.0001) {
    Output = 'Reject'
  }else if(MW_p <= 0.0001 & KS_p > 0.0001) {
    Output = 'MW Reject'
  }else{
    Output = 'Reject'
  }
  return(paste(Output, sep = ' '))
}
WorkingMatrix = matrix(nrow = 4, ncol = 4)
WorkingMatrixColours = matrix(nrow = 4, ncol = 4)
WorkingMatrix[1,1] = NA
WorkingMatrix[1,2] = StatisticalDifferenceTest(ALSC011_Central_Normal_S1_Sample_29, ALSC011_Central_6m_S1_Sample_29)
WorkingMatrix[1,3] = StatisticalDifferenceTest(ALSC011_Central_Normal_S1_Sample_29, ALSC011_Central_12m_S1_Sample_29)
WorkingMatrix[1,4] = StatisticalDifferenceTest(ALSC011_Central_Normal_S1_Sample_29, ALSC011_Central_24m_S2_Sample_29)
WorkingMatrix[2,1] = StatisticalDifferenceTest(ALSC011_Central_Normal_S1_Sample_29, ALSC011_Central_6m_S1_Sample_29)
WorkingMatrix[2,2] = NA
WorkingMatrix[2,3] = StatisticalDifferenceTest(ALSC011_Central_6m_S1_Sample_29, ALSC011_Central_12m_S1_Sample_29)
WorkingMatrix[2,4] = StatisticalDifferenceTest(ALSC011_Central_6m_S1_Sample_29, ALSC011_Central_24m_S2_Sample_29)
WorkingMatrix[3,1] = StatisticalDifferenceTest(ALSC011_Central_Normal_S1_Sample_29, ALSC011_Central_12m_S1_Sample_29)
WorkingMatrix[3,2] = StatisticalDifferenceTest(ALSC011_Central_6m_S1_Sample_29, ALSC011_Central_12m_S1_Sample_29)
WorkingMatrix[3,3] = NA
WorkingMatrix[3,4] = StatisticalDifferenceTest(ALSC011_Central_24m_S2_Sample_29, ALSC011_Central_12m_S1_Sample_29)
WorkingMatrix[4,1] = StatisticalDifferenceTest(ALSC011_Central_Normal_S1_Sample_29, ALSC011_Central_24m_S2_Sample_29)
WorkingMatrix[4,2] = StatisticalDifferenceTest(ALSC011_Central_6m_S1_Sample_29, ALSC011_Central_24m_S2_Sample_29)
WorkingMatrix[4,3] = StatisticalDifferenceTest(ALSC011_Central_12m_S1_Sample_29, ALSC011_Central_24m_S2_Sample_29)
WorkingMatrix[4,4] = NA
for(m in 1:nrow(WorkingMatrix)){
  for(o in 1:ncol(WorkingMatrix)){
    if(is.na(WorkingMatrix[m,o])){
      WorkingMatrixColours[m,o]= 'grey'
    }else if(WorkingMatrix[m,o] == 'Reject'){
      WorkingMatrixColours[m,o]= 'red'
    }else if(WorkingMatrix[m,o] == 'Accept'){
      WorkingMatrixColours[m,o]= 'green'
    }else{
      WorkingMatrixColours[m,o]= 'yellow'
    }
  }
}
WorkingMatrix
```
